# Exploring Protein-DNA Binding Residue Prediction and Consistent Interpretability Analysis Using Deep Learning

**DOI:** 10.1101/2024.10.12.613667

**Authors:** Yufan Liu

## Abstract

Accurately identifying DNA-binding residues is a crucial step in developing computational tools to model DNA-protein binding properties, which is essential for binding pocket discovery and related drug design. Although several tools have been developed to predict DNA-binding residues based on protein sequences and structures, their performance remains limited, and proteins with crystal structures still represent only a small fraction of DNA-binding proteins. Additionally, the process of extracting handcrafted features for protein representation is labor-intensive. In this study, we combined the strengths of pre-trained protein language models and attention mechanisms to propose a sequence-based method: an attention-based deep learning approach for accurately predicting DNA-binding residues, incorporating a contrastive learning module. Our method outperformed all other sequence-based models across two prevalent benchmark datasets. Furthermore, we developed a structure-based graph neural network (GNN) model to demonstrate the impact of the contrastive module. A common limitation of existing models is their lack of interpretability, which hinders our ability to understand what these models have learned. To address this, we introduced a novel perspective for interpreting our sequence-based model by analyzing the consistency between attention scores and the edge weights generated by the GNN model. Interestingly, our results show that large-scale pre-trained protein language models, together with attention mechanisms, can effectively capture structural information solely from protein sequence inputs.

## 1 Introduction

The interaction between proteins and ligands plays a pivotal role in nearly all biological activities within living organisms. These interactions encompass protein-protein, protein-molecule, and protein-nucleic acid interactions. Of particular significance is DNA, which, as the carrier of genetic instructions, interacts with proteins to facilitate crucial biological processes such as transcription, replication, gene expression, signal transduction, and metabolism [1, 2, 3].

Over the past few decades, a multitude of computational approaches have been proposed for predicting DNA-binding sites. These methods can be broadly categorized into two types: sequence-based and structure-based. Sequence-based methods include tools such as DRNAPred [4], DNAPred [5], SVMnuc [6], NCBRPred [7] and DBpred [8], while structure-based methods include models like DNABind [9], COACH-D [10], NucBind [6] GraphBind [11], and GraphSite [12]. For the purposes of this discussion, structure-based models are narrowly defined as those that utilize the spatial coordinates of proteins, whereas models that rely on predefined secondary structural information are classified as sequence-based.

Sequence-based methods utilize features extracted directly from protein sequences, such as one-hot encoding and relative positional representations. Recently, the DBPred method introduced a novel sequence-based feature that employs a 17-mer pattern. Generally, these models also incorporate evolutionary information by leveraging features derived from multiple protein sequences. Common tools used for generating such evolutionary information include PSI-BLAST [13] and HHblits [14], which produce multiple sequence alignments (MSAs) and position-specific scoring matrices (PSSMs). Extensive research has demonstrated that evolutionary information significantly improves performance in ligand-binding prediction tasks [11, 8]. Studies have further shown that protein-DNA binding preferences are partially dictated by structural and physicochemical characteristics from both the protein and DNA, known as shape readout and base readout, respectively [15]. To enhance prediction outcomes, structural information is often integrated, such as that provided by DSSP [16, 17] for secondary structure assignment, while physicochemical properties like hydrophobicity and polarity are also recognized as effective features [18].

In contrast, structure-based models have developed rapidly, driven by the increasing availability of protein-DNA complex structures in the Protein Data Bank [19]. The subdivision of structure-based models has been comprehensively categorized in previous studies [12]. Template-based models, such as COACH-D, first generate a predictive structure using a sequence template, followed by a docking process to identify potential DNA-binding pockets. This approach fully leverages structural information in a physical manner, despite being time- and resource-intensive. The development of deep learning techniques and their application to graph-structured data has led to the emergence of several graph neural network (GNN)-based models, such as GraphBind and GraphSite. GNNs are widely applied to biological data due to their natural graph structure. To capitalize on coordinate information, models such as S-MAN [20] and GeoPPI [21] have been proposed for various biological prediction tasks. These DNA-binding site prediction models utilize the GNN’s ability to aggregate information from connected neighborhoods, achieving high performance on benchmark datasets. For example, the GraphSite model uses a graph Transformer with K-nearest neighbors to incorporate structural information. Notably, all structure-based models can also utilize sequence-based features as input representations; for instance, GraphBind incorporates both structural information and evolutionary matrices as initial features. As a result, structure-based models generally outperform sequence-based models. However, despite their strong performance, the effectiveness of structure-based models is limited by the scarcity of available protein-DNA complex structures, and many DNA-binding protein sequences lack accurate crystal structures. Thus, the development of robust and accurate sequence-based models for predicting DNA-binding residues remains a critical challenge.

In this study, we developed an attention-based deep learning model to predict DNA-binding residues from protein sequences, leveraging the pre-trained protein language model and the interpretability provided by the multi-head attention mechanism. We incorporated a contrastive learning module into the loss computation process, which enhances model performance by learning discriminative representations in the embedding space. Furthermore, we introduced a simple structure-based model using a graph neural network to verify that the contrastive learning module is effective for both sequence-based and structure-based models, demonstrating its potential as a general component in ligand-binding residue prediction frameworks. Finally, we systematically analyzed the interpretability of both sequence-based and structure-based models, revealing underlying consistencies between them. This analysis improves the understanding of model design for ligand-binding residue identification tasks.

## 2 Materials and methods

### 2.1 Datasets

We selected two widely used benchmark datasets to evaluate and compare the performance of our proposed model against existing methods. For clarity, we refer to these datasets as Dataset1 and Dataset2, and denote the training and testing subsets as TR and TE, respectively. Notably, both datasets were pre-processed using a similar procedure to enhance model robustness and mitigate bias from imbalanced data distribution. This included reducing sequence similarity using a 30

#### Dataset1 Description

This dataset was originally created for the DBPred study, a sequence-based deep learning method for predicting DNA-binding residues in proteins [8]. It was compiled from the hybridNAP [22] and ProNA2020 [23] databases. The training set (denoted as TR646) contains 646 proteins with 15,636 DNA-binding residues and 298,503 non-binding residues, while the testing set (denoted as TE646) includes 46 proteins with 956 DNA-binding residues and 9,911 non-binding residues.

#### Dataset2 Description

This dataset was first proposed in the GraphBind study, which developed a structure-based graph neural network (GNN) model for recognizing nucleic acid-binding residues. It was subsequently used by GraphSite, another GNN model designed to predict DNA-binding residues based on protein structures generated by AlphaFold2. Dataset2 consists of a training set (denoted as TR573) of 573 proteins with 14,479 DNA-binding residues and 145,404 non-binding residues, and a testing set (denoted as TE129) of 129 proteins with 2,240 DNA-binding residues and 35,275 non-binding residues. To address data imbalance, GraphBind applied data augmentation to the training set, and we followed the same augmentation approach in this study.

Importantly, Dataset1 serves as a benchmark for sequence-based models, while Dataset2 is a benchmark for structure-based models. Thus, we evaluated and compared our sequence-based model on both Dataset1 and Dataset2, but restricted the evaluation of our structure-based model to Dataset2. The protein structures in Dataset2 were obtained from the BioLiP database [24] using the corresponding PDB accession numbers. To ensure fairness in comparisons, we adhered to the same data-splitting strategies as originally proposed for both datasets.

### 2.2 Architecture of the Proposed Model

The overall architecture of the proposed model is illustrated in Figure 1. As depicted, the model is divided into two primary components: a sequence-based model (Figure 1A, B, C, and D) and a structure-based model (Figure 1D, E, and F). Both models employ a pre-trained protein language model as a feature extractor, without any fine-tuning, and a contrastive learning (CL) model to enhance protein representation in the embedding space. The details of these components will be elaborated in subsequent sections. The sequence-based model was used to evaluate and compare the prediction performance on protein sequences, while the structure-based model was employed to analyze the contributions of the sequence-based components and provide interpretability for DNA-binding residue predictions.

**Figure 1:**
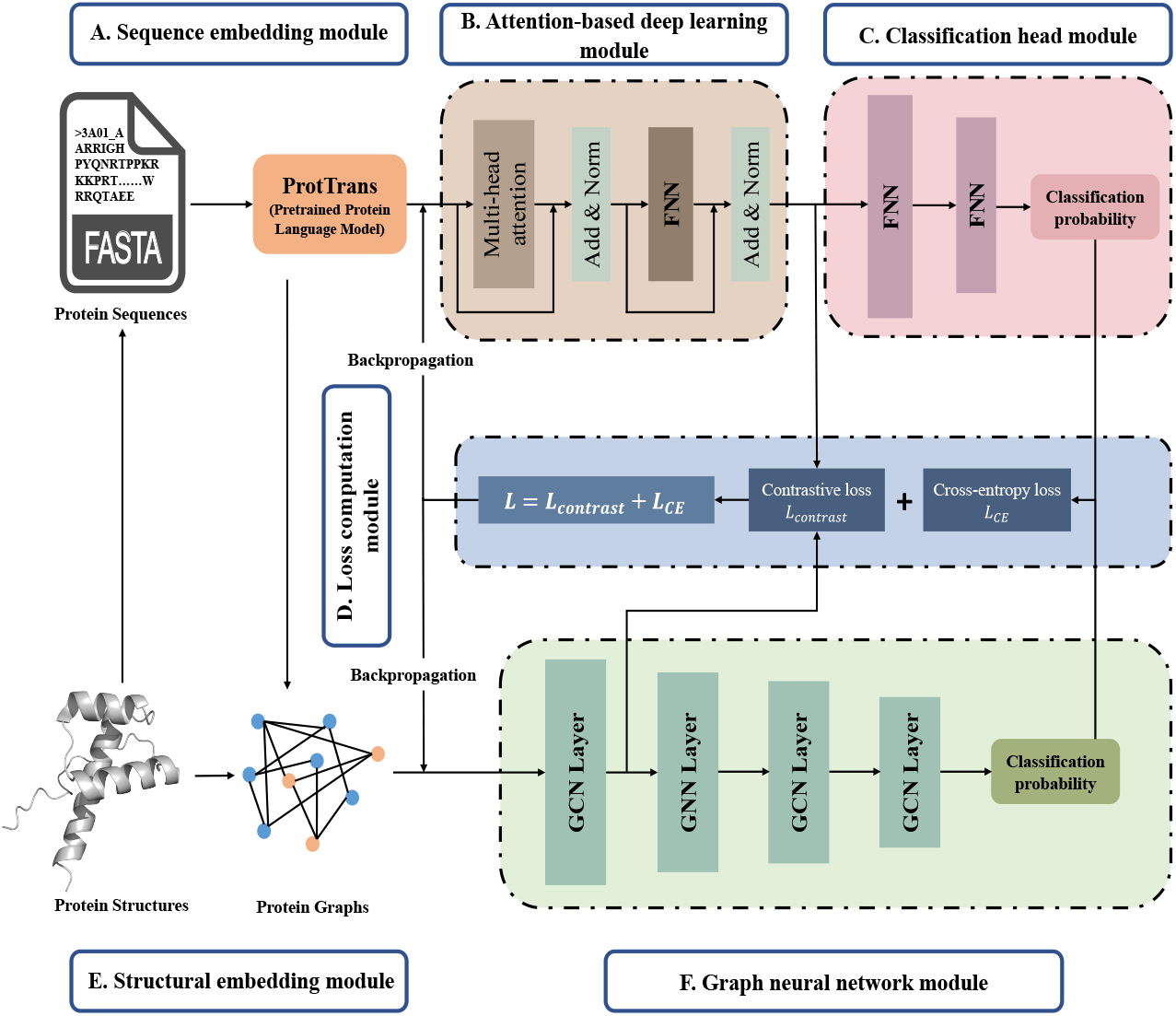
An overall architecture of our proposed sequence-based model and the parallel structure-based model. (A) The sequence embedding module generates protein sequence representations using a pre-trained protein language model, ProtBert. (B) The attention-based deep learning model captures long-range dependencies within the protein sequence, enhancing interpretability. (C) The classification head module outputs the classification probability for each residue. (D) The loss computation module calculates cross-entropy and contrastive loss, which are used in backpropagation to update model parameters. (E) The structure construction model builds a topological graph of protein structures to facilitate structure-based model learning. (F) The graph neural network module evaluates the contributions of components from the sequence-based model and provides interpretability from a structural perspective.

#### 2.2.1 Protein Sequence Embedding Module

Raw protein sequences were processed using a large-scale pre-trained language model, ProtTrans [25], to obtain protein embeddings. ProtTrans is a set of models pre-trained on vast amounts of protein sequence data and is widely used in protein structure and property prediction tasks. Specifically, we adopted ProtBert, a BERT-based model with approximately 400 million parameters, as the feature extractor. BERT (Bidirectional Encoder Representations from Transformers) [26, 27] uses a masked language model strategy to embed tokens with context from both directions, making it highly effective for token-level tasks such as sentiment analysis and named entity recognition. Given that DNA-binding residue prediction is also a token-level classification task, ProtBert was used without fine-tuning during the training process to obtain initial protein embeddings. The same feature extraction method was employed for node features in the structure-based model, and the sequence embedding process was carried out using the HuggingFace Transformers library [28].

#### 2.2.2 Attention-Based Deep Learning Module

To capture long-range interactions within protein sequences and improve interpretability, we employed an attention-based deep learning module as the encoder. This module consists of a multi-head attention block, a fully connected neural network (FNN), and residual connections. The multi-head attention mechanism allows the model to jointly learn different subparts of the input protein embeddings using parallel self-attention modules. The self-attention mechanism is defined as follows:

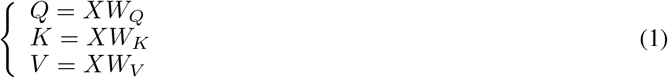

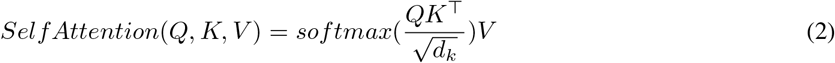

Here, 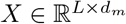 represents the protein embedding generated by the protein sequence embedding module, which is then transformed into the query matrix 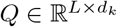, key matrix 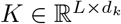, and value matrix 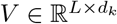 through linear transformations. Given the embeddings from ProtBert, *d*_*m*_ is set to 1024. The multi-head attention mechanism is formulated as follows:

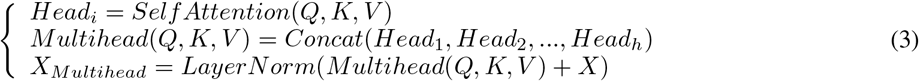

Here, 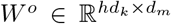 is a linear transformation matrix that restores the attention dimension to the original embedding dimension. The number of attention heads is denoted by *h* and we set *hd*_*k*_ = *dm* for computational convenience. Residual connections and layer normalization were applied to enhance performance, following the post-norm approach based on experimental results. The final component of the module is an FNN layer, defined as follows:

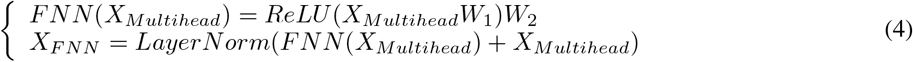

Here, 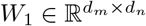 and 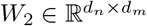 are linear transformation matrices, and *d*_*n*_ is the hidden dimension of the FNN layer. A rectified linear unit (ReLU) is used as the activation function to introduce nonlinearity, and the same residual and normalization techniques are applied. The output of this module is fed into the classification and loss computation modules for further processing.

#### 2.2.3 Classification Head Module

The sequence representation generated by the attention-based module is passed into the classification head module to compute the probabilities for the negative (non-binding) and positive (DNA-binding) classes. The classification head consists of multiple FNN layers, described mathematically as follows:

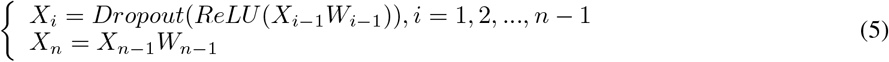

Here, *X*_0_ is the output sequence representation from the previous module. ReLU is applied as the activation function, and the dropout technique is employed to stabilize training. Dropout and ReLU are applied to all layers except the final one, where a simple linear transformation produces the output.

#### 2.2.4 Structure Construction Module

To analyze the contribution of sequence-based components and interpret the predicted DNA-binding residues, we designed a simple graph neural network (GNN) model in parallel. In the structure embedding module, proteins are represented as graphs where each residue serves as a node. The protein structure is represented as a topological graph *G* = (*V, E*, ***A***), where *V, E*, ***A*** represent the set of vertices, edges, and adjacency matrix, respectively.

We used the same pre-trained protein embeddings as node features (Figure 1A), with a dimension of 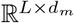, where *d*_*m*_ is set to 1024. For edges, we established a distance-based cutoff: if the distance between two nodes is less than the cutoff, an edge is created. The distance between nodes is defined as the Euclidean distance between the C_*α*_ atoms of the corresponding residues. Edge weights were set as the reciprocal of this distance, with self-loop edge weights set to 1. The graph construction process is mathematically expressed as:

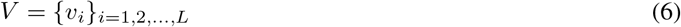

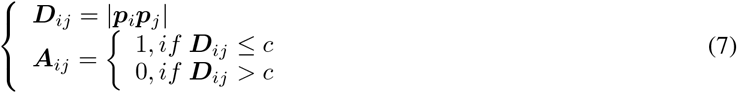

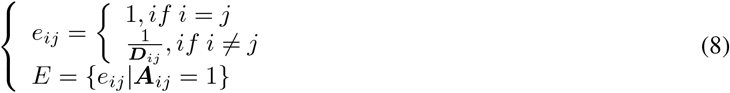

Here, *v*_*i*_ represents node features generated by ProtBert. ***D***_*ij*_ represents the distance matrix, with *p*_*i*_ denoting the Cartesian coordinates of the *i*-th residue. Edges were formed based on a distance threshold *c*, and edge weights were computed using Equation 8.

#### 2.2.5 Graph Neural Network Module

Graph neural networks (GNNs) excel in capturing topological relationships in networks such as social, transportation, molecular, and protein interaction networks. Most GNN architectures fall under the message-passing neural network (MPNN) framework [29]. In this study, we employed a GNN to evaluate the composition of the sequence-based model and provide interpretability. Specifically, we used a stack of graph convolutional network (GCN) layers [30]. The GCN is formulated as follows:

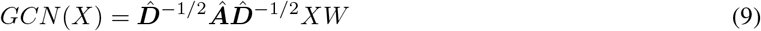

Here, ***Â*** = ***A*** + ***I*** is the adjacency matrix with a self-loop, and 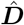 denotes the degree diagonal matrix. 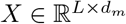 is the node embedding in each GCN layer, and *W ∈ d*_*m*_*× d*_*k*_ is the parameter matrix that transforms the node features to the output dimension *d*_*k*_. We applied batch normalization, dropout, and ReLU activation across all GCN layers except the last one. A classification head module was used to generate classification probabilities as described above. Importantly, we applied the same contrastive loss module to both sequence-based and structure-based models.

To maintain consistency, the first GCN layer’s transformation matrix was set to 1024× 1024, consistent with the attention-based module’s output, which was then passed to the loss computation module.

#### 2.2.6 Loss Computation Module

The loss between the predicted values and the ground truth labels is computed by the loss computation module. This module consists of two main components: cross-entropy loss and contrastive learning loss. The inputs for these two components are derived from an intermediate embedding layer and the classification head module, respectively.

##### Contrastive Learning Loss

The objective of contrastive learning is to map similar samples close to each other in the embedding space, while pushing dissimilar samples further apart. Contrastive learning techniques are widely adopted in fields such as computer vision and natural language processing, with models like SimCLR [31] and MOCO [32] demonstrating strong capabilities in representation learning. In protein-ligand binding prediction tasks, contrastive learning has been shown to improve model performance, particularly in scenarios involving imbalanced datasets [33]. Building on these insights, we integrated a contrastive learning module into both the sequence-based and structure-based models.

To compute the contrastive loss, we first obtained a batch of protein representations with a dimension of 1024 from the attention-based deep learning module. The contrastive loss is computed by comparing the first half of the batch with the second half. It is important to note that, because our model does not require fine-tuning of the pre-trained model, we can increase the batch size without additional computational overhead. This larger batch size provides more positive and negative residue embeddings for contrastive loss computation, which in turn enhances model performance. The contrastive learning loss is defined as follows:

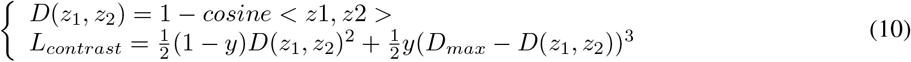

Here, *z*_1_ and *z*_2_ represent arbitrary embeddings of residues, and the distance between them is measured using cosine similarity. is to maximize the distance between the embeddings of DNA-binding residues, while minimizing the distance between DNA-binding residues and non-binding residues. The variable *y* is assigned a value of 0 when the two residues are either both DNA-binding or both non-binding, and 1 when the two residues belong to different classes. The maximum distance value, *D*_*max*_ is set to 2, and a higher penalty is applied to the loss between different classes to account for the imbalance in the dataset distribution.

##### Cross-Entropy Loss

Cross-entropy loss (denoted as CE in the formulation) is a commonly used objective function for classification tasks. It can be formulated as:

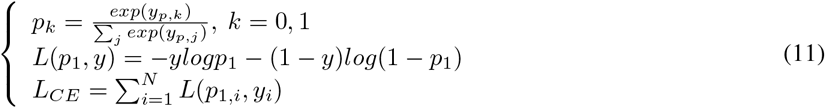

Here, *k* represents the different classes, positive or negative, and *p* denotes the classification probability for class *k. N* is the number of residues in a single batch, and *y*_*i*_ is the true label of the target residue. In summary, the overall loss of our model is calculated as *L* = *L*_*contrast*_ + *L*_*CE*_. Although the contrastive learning module is designed to address data imbalance, we applied a weighted cross-entropy loss during training to penalize incorrectly classified DNA-binding residues. It is important to note that backpropagation is halted at the embedding layer generated by ProtBert, meaning the pre-trained model remains unaltered.

### 2.3 Evaluation Metrics

To maintain consistency with previous studies [11, 12, 8], we utilized several classification evaluation metrics, including specificity (Spe), precision (Pre), recall (Rec), F1-score (F1), Matthews correlation coefficient (MCC), and the area under the receiver operating characteristic curve (AUC). These metrics are defined as follows:

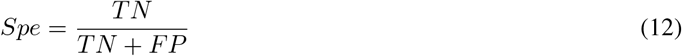

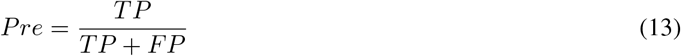

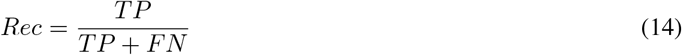

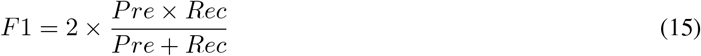

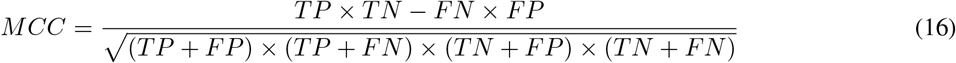

Here, TP, FP, TN, and FN represent true positives (the number of residues correctly classified as DNA-binding sites), false positives (the number of residues incorrectly classified as DNA-binding sites), true negatives (the number of residues correctly classified as non-binding sites), and false negatives (the number of residues incorrectly classified as non-binding sites), respectively. Specifically, specificity (Spe) indicates the proportion of correctly predicted non-binding sites, while precision (Pre) measures the accuracy of residues predicted to be DNA-binding sites. Recall (Rec) assesses the model’s ability to correctly identify DNA-binding residues, and the F1-score (F1) is the harmonic mean of precision and recall. The Matthews correlation coefficient (MCC) evaluates the model’s overall prediction performance across both classes (binding and non-binding), making it particularly useful for imbalanced datasets. Additionally, the area under the receiver operating characteristic curve (AUC) provides a measure of the model’s overall performance across various classification thresholds.

## 3 Results

### 3.1 Performance Evaluation of the Sequence-Based Model

We first compared the performance of our sequence-based model to existing models using the evaluation metrics described in the previous section. The model was tested on both Dataset1 and Dataset2, ensuring a fair comparison by training TE46 only on TR646 and TE129 only on TR573. We evaluated our model against sequence-based models on Dataset1 (Table 2), and both sequence-based and structure-based models on Dataset2 (Table 3). It is important to note that NucBind is an ensemble model that incorporates predictions from both COACH-D and SVMnuc. While COACH-D uses protein structure predictions, SVMnuc only uses sequence-based features, but since NucBind accepts only protein sequences as input, it can be categorized as both a sequence- and structure-based model. In Table 3, we have classified NucBind as a structure-based model.

**Table 1:**
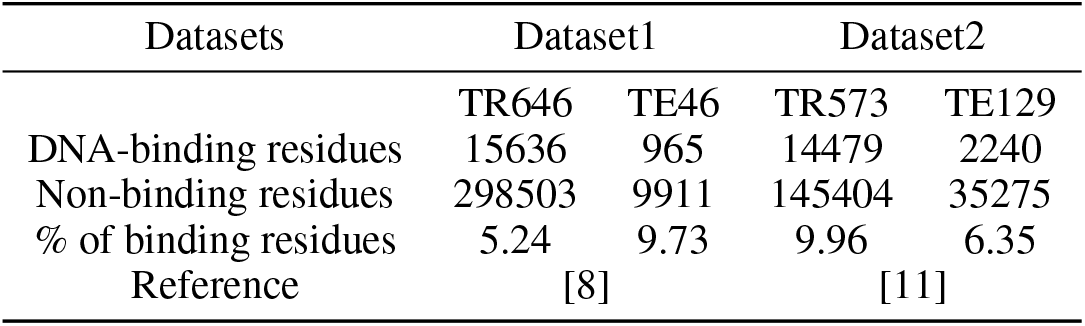
Summary of benchmark datasets.

**Table 2:**
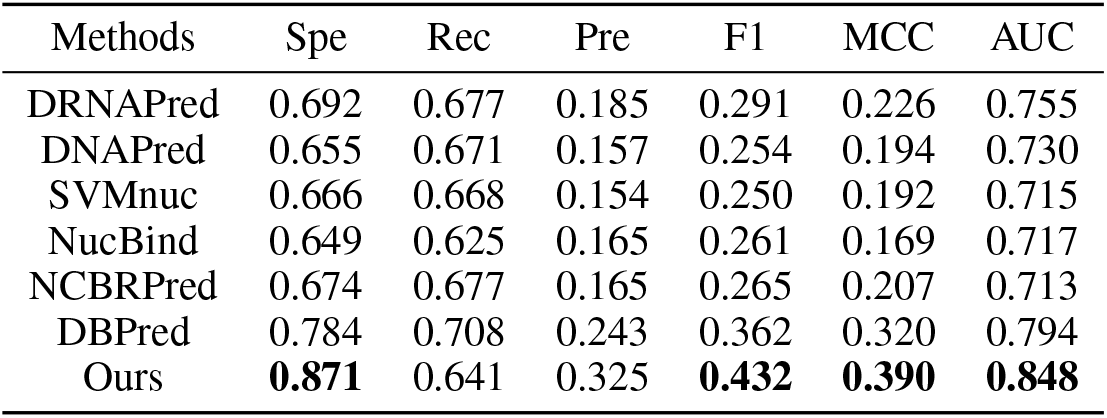
Comparison of sequence-based model with other methods on TE46.

| Methods | Spe | Rec | Pre | F1 | MCC | AUC |
| --- | --- | --- | --- | --- | --- | --- |
| DRNAPred | 0.692 | 0.677 | 0.185 | 0.291 | 0.226 | 0.755 |
| DNAPred | 0.655 | 0.671 | 0.157 | 0.254 | 0.194 | 0.730 |
| SVMnuc | 0.666 | 0.668 | 0.154 | 0.250 | 0.192 | 0.715 |
| NucBind | 0.649 | 0.625 | 0.165 | 0.261 | 0.169 | 0.717 |
| NCBRPred | 0.674 | 0.677 | 0.165 | 0.265 | 0.207 | 0.713 |
| DBPred | 0.784 | 0.708 | 0.243 | 0.362 | 0.320 | 0.794 |
| Ours | **0.871** | 0.641 | 0.325 | **0.432** | **0.390** | **0.848** |

**Table 3:** Comparison of sequence-based model with other methods on TE129.

| Method | Spe | Rec | Pre | F1 | MCC | AUC |
| --- | --- | --- | --- | --- | --- | --- |
| DRNAPred | 0.937 | 0.233 | 0.190 | 0.210 | 0.155 | 0.693 |
| DNAPred | 0.954 | 0.396 | 0.353 | 0.373 | 0.332 | 0.845 |
| SVMnuc | 0.966 | 0.316 | 0.371 | 0.341 | 0.304 | 0.812 |
| NCBRPred | 0.969 | 0.312 | 0.392 | 0.347 | 0.313 | 0.823 |
| DBPred | 0.958 | 0.070 | 0.100 | 0.080 | 0.032 | 0.526 |
| COACH-D <sup>a</sup> | 0.958 | 0.367 | 0.357 | 0.362 | 0.321 | 0.710 |
| NucBind <sup>a</sup> | 0.966 | 0.330 | 0.381 | 0.354 | 0.317 | 0.811 |
| DNABind <sup>a</sup> | 0.926 | 0.601 | 0.346 | 0.440 | 0.411 | 0.858 |
| GraphBind<sup>a</sup> | 0.941 | 0.684 | 0.422 | 0.522 | 0.500 | 0.928 |
| GraphSite <sup>a</sup> | 0.950 | 0.665 | 0.460 | 0.543 | 0.519 | 0.934 |
| Ours (seq) <sup>b</sup> | 0.831 | 0.717 | 0.213 | 0.328 | 0.324 | 0.867 |
| Ours (struct) <sup>c</sup> | 0.862 | 0.681 | 0.239 | 0.353 | 0.342 | 0.864 |

<sup>a</sup> refers to structure-based models
<sup>b</sup> refers to a sequence-based model
<sup>c</sup> refers to a structure-based model

As shown in Table 2, our model significantly outperformed all sequence-based models on Dataset1 and some structure-based models on Dataset2. Specifically, on Dataset1, our model achieved a specificity of 0.871, a precision of 0.325, an F1-score of 0.432, an MCC of 0.390, and an AUC of 0.848. This represents a substantial improvement over the second-best model, DBPred, with gains of 11.1

On Dataset2 (Table 3), our model outperformed all sequence-based models and three structure-based models (COACH-D, NucBind, and DNABind) on the AUC metric, and surpassed two structure-based models on the MCC metric. Notably, the second-best sequence-based model achieved an AUC of 0.526, only slightly above the random classification threshold (0.5). To assess the generalization capability of our model, we transferred the model weights trained on TR646 to TE129, following the same protocol used for DBPred on Dataset2. As shown in Figure 2A, our model trained on TR646 also demonstrated strong performance, with an AUC of 0.855, which is comparable to the performance of the model trained on TR573. The ROC curve of the transferred model closely matched that of the model trained on TR573, indicating robust generalization to unseen datasets, a capability that the DBPred model lacks. We argue that models should generalize well on datasets with similar distributions, such as DNA-binding protein sequences, and our model demonstrated this principle effectively in the experiments.

**Figure 2:**
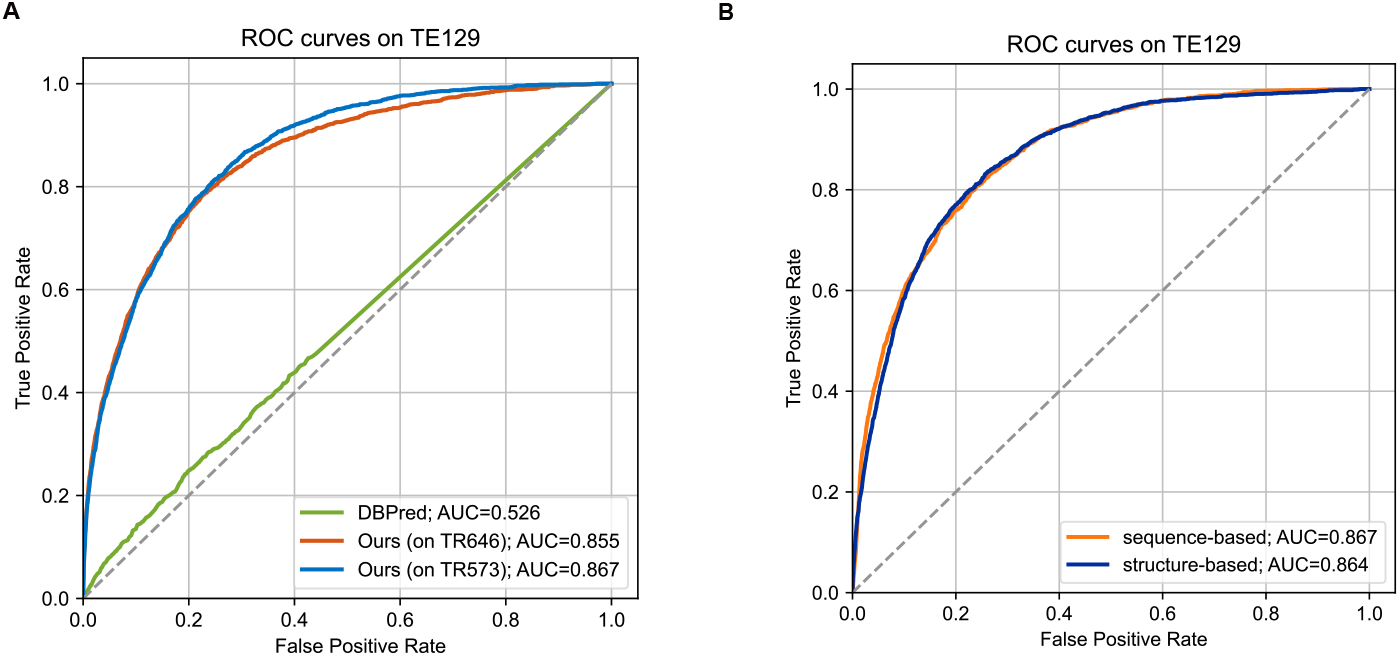
ROC curves comparing our proposed model with existing methods: (A) ROC curves for DBPred and our model on the TE129 dataset, where the red and blue lines represent the performance of our model trained on TR646 and TR573, respectively, while the green line shows DBPred’s performance and the dashed line indicates random classification. (B) ROC curves comparing the sequence-based model (orange) and structure-based model (blue) trained on TR573.

We also evaluated the performance of the parallel structure-based model, which consists of a stack of GCN layers. The ROC curves for both structure-based and sequence-based models are shown in Figure 2B. Interestingly, the sequence-based model exhibited competitive performance compared to the structure-based model, with nearly identical ROC curves. This suggests that the sequence-based model may have captured similar structural information as the structure-based model. Since both models utilized the same pre-trained ProtBert model, this result can be interpreted through the lens of the global attention mechanism. In the attention-based deep learning model, the attention block captures long-range token dependencies within protein sequences, a process analogous to the graph construction procedure in the structure-based model. As a result, it is unsurprising that the attention mechanism produces similar outcomes to a GNN-based model. Additionally, in structure-based models, an accurate spatial representation of the protein enhances model performance, which may explain why our sequence-based model outperformed some structure-based models that relied on predicted structures, such as COACH-D. However, with advancements in protein structure prediction, such as the development of AlphaFold2 [34], the accuracy of predicted structures has improved dramatically, leading to the emergence of AlphaFold2-based models, such as GraphSite, which currently achieves the highest performance among structure-based models.

In conclusion, as evidenced by the results in Table 2, Table 3, and Figure 2, our sequence-based model achieved state-of-the-art performance on both Dataset1 and Dataset2, surpassing several structure-based models. Additionally, our model demonstrated better generalization capability than the recently proposed DBPred model, indicating its robustness. Moreover, the sequence-based model was able to learn representations in a manner comparable to structure-based models, largely due to the global attention mechanism.

### 3.2 The Contrastive Learning Module Improves the Performance of Both Sequence-Based and Structure-Based Models

To assess the effectiveness of the contrastive learning module in both sequence- and structure-based models, we re-trained the models without the contrastive learning component. For consistent comparison, both models were evaluated on Dataset2, and the performance results are presented in Table 4. As indicated in the table, models incorporating the contrastive learning module outperformed those without it in terms of specificity, F1-score, MCC, and AUC. This demonstrates that the contrastive learning loss enhances the model’s capacity to distinguish between different residue classes in the embedding space. Among these metrics, specificity shows that the contrastive learning module improved the identification accuracy of non-binding residues, reducing the false negative rate. Given that MCC is particularly sensitive to imbalanced data distribution, the contrastive learning module also proves beneficial in improving model performance under such conditions, as evidenced by our experimental results.

**Table 4:** Evaluation of the effectiveness of contrastive learning module on TE129.

| Method | CL module | Spe | Rec | Pre | F1 | MCC | AUC |
| --- | --- | --- | --- | --- | --- | --- | --- |
| sequence-based | - | 0.819 | 0.710 | 0.199 | 0.312 | 0.307 | 0.861 |
|  | + | 0.831 | 0.717 | 0.213 | 0.328 | 0.324 | 0.867 |
| structure-based | - | 0.849 | 0.686 | 0.224 | 0.337 | 0.328 | 0.853 |
|  | + | 0.862 | 0.681 | 0.239 | 0.353 | 0.342 | 0.864 |

To further validate that the contrastive learning module facilitates a more discriminative embedding space, we conducted a t-Distributed Stochastic Neighbor Embedding (t-SNE) analysis [35] on the embeddings used for contrastive loss computation. The embeddings had a dimensionality of 1024, as previously described. The results of the t-SNE analysis are depicted in Figure 3. The embedding space produced by the model with the contrastive learning module is more distinguishable, particularly in the case of the structure-based model. The contrastive learning module’s impact on the sequence-based model is comparatively limited, potentially due to the attention-based deep learning module already capturing sufficient information for classification. Consequently, the contrastive learning module provides only incremental improvement.

**Figure 3:**
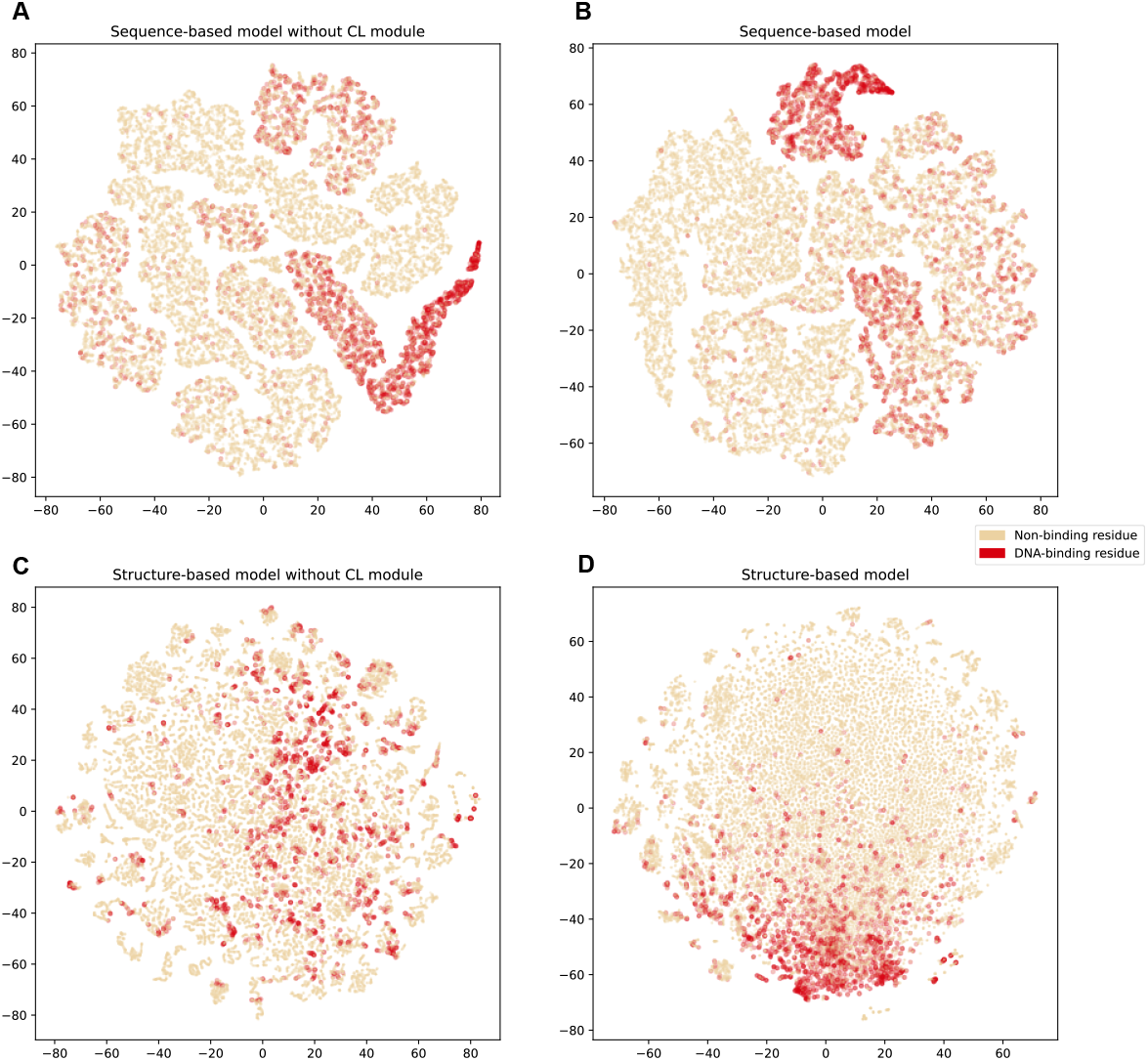
The t-SNE reduction visualization of embedding space of our sequence-based and structure-based models tested on TE129. (A, B) The embedding space of sequence-based model without and with the contrastive learning module. (C, D) The embedding space structure-based model without and with the contrastive learning module.

To test this assumption, we replaced the attention-based module with a simple multi-layer perceptron (MLP) and visualized the reduced embeddings (of dimension 1024) using t-SNE. As shown in Supplementary Figure 1, the model with the contrastive learning module clearly produced more discriminative embeddings, indicating that contrastive learning is particularly beneficial when applied to shallower network layers. This highlights the importance of the module’s placement within the model architecture. Moreover, both models with and without the contrastive learning module learned more discriminative embeddings compared to the initial ProtBert-generated embeddings, as shown in Supplementary Figure 2.

Additionally, we explored the effect of embedding dimension size on model performance, as illustrated in Figure 4. Using AUC and MCC as evaluation metrics, we tested various embedding dimensions in the structure-based model on TR573. Notably, an embedding dimension of 2 means that contrastive loss was computed directly using the classification probabilities from the classification head module. As shown in the figure, the dimension size of 1024 yielded the best performance, while smaller dimensions, such as 2, resulted in worse AUC and MCC scores compared to models without the contrastive learning module. This underscores the importance of selecting an appropriate embedding dimension to maximize the effectiveness of contrastive learning, with larger dimensions leading to more discriminative representations.

**Figure 4:**
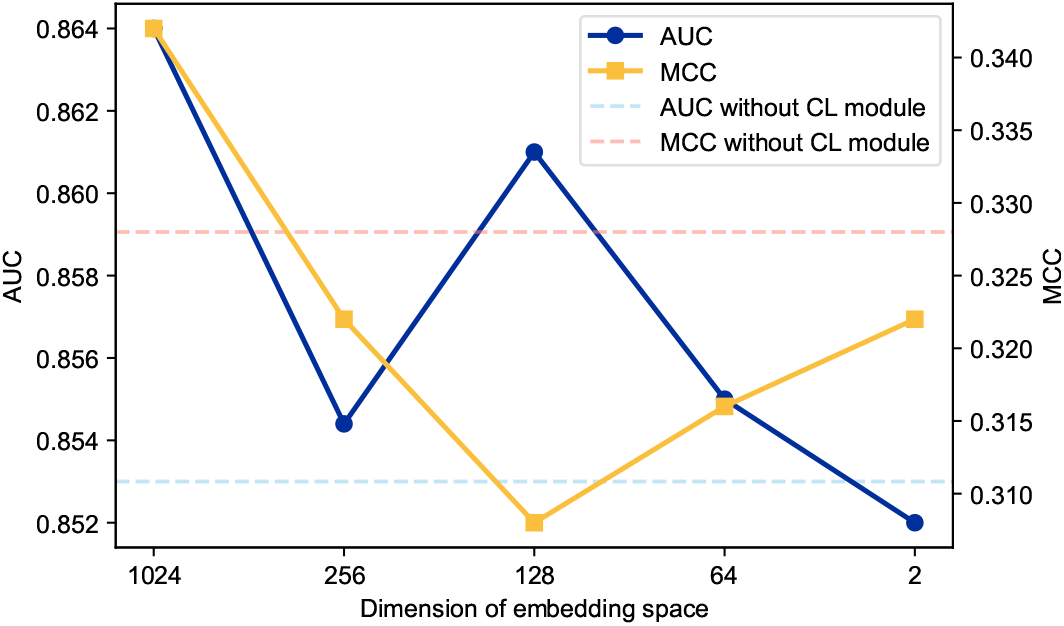
The correlation between the dimension of the embedding space and the AUC, as well as the MCC, of the structure-based model is depicted. The navy blue line represents the changes in AUC with varying embedding dimensions, while the yellow line represents the MCC. The two dashed lines indicate the AUC and MCC values for models without the contrastive learning module, with the sky blue line showing AUC and the purple line showing MCC.

In summary, the contrastive learning module effectively improves performance in both sequence-based and structure-based models and enhances the distinction between residue classes in various embedding dimensions. The module’s placement within the model architecture and the size of the embedding space are critical factors that must be determined through experimentation. In line with previous studies [33], contrastive learning can be considered a flexible and essential component for ligand-binding residue identification frameworks, contributing to improved model performance.

### 3.3 The Sequence-Based Model Learned Structural Information via Model Interpretability Analysis

The advancement of deep neural networks has significantly improved the accuracy of DNA-binding residue prediction; however, many of these models operate as black boxes with limited interpretability. Without a clear understanding of the underlying mechanisms, such models cannot be fully trusted. Additionally, the field of DNA-binding residue identification has seen a lack of systematic interpretability analysis. In this study, we conducted an interpretability analysis of our sequence-based and structure-based models in parallel, aiming to explore the contribution of neighboring residues to the accurate prediction of DNA-binding residues from both sequence and structural perspectives.

As shown in the overall model architecture in Figure 1, we did not fine-tune the pre-trained model, but the attention-based deep learning module inherently provided interpretability through attention scores [36]. In contrast, graph neural networks (GNNs) have been the subject of interpretability research, using various strategies such as perturbation-based approaches (e.g., GNNExplainer [37] and PGExplainer [38]) and surrogate-based models (e.g., GraphLime [39] and PGM-Explainer [40]). These models demonstrate interpretability from a *post-hoc* perspective, which helps enhance understanding of GNN-based models.

To fairly compare the interpretability of our sequence-based and structure-based models, we first evaluated their performance on TE129. Supplementary Figure 3 illustrates the correlation of AUC values per sequence, showing a high degree of relevance between the two models, particularly when predictions are highly accurate. We selected proteins with minimal prediction differences (less than 0.010) and high accuracy (sequence-based model AUC greater than 0.950) for further interpretability analysis. The PDB accession numbers and predicted AUC values for these proteins are listed in Supplementary Table 1.

At the sequence-based level, we visualized the attention weights of target binding residues to analyze the contributions of other residues. At the structure-based level, we utilized a modified version of PGExplainer [38] to assess the contribution of neighboring nodes to a given DNA-binding site. PGExplainer is a parameterized explainer for GNNs that identifies a key subgraph *G*_*s*_ which maximizes the mutual information between the subgraph and the original graph. In the original version of PGExplainer, graphs are constructed without edge weights, and the generated edge weights represent the importance of masked edges. However, since edge weights in our study represent the reciprocal of the distance between residues, we made a slight modification to the importance computation: masked edge weights were multiplied by their original values to model edge importance. The remaining mutual information maximization procedure remained the same as in PGExplainer. The re-weighted edge attribute can be mathematically described as follows:

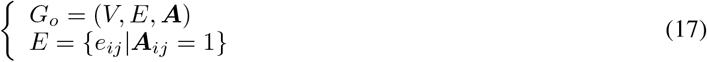

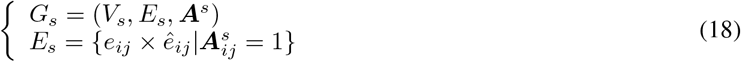

Here, *G*_*o*_ represents the original graph and *G*_*s*_ denotes the sampled subgraph. It is important to note that *e*_*ij*_ is the edge weight in the original graph, as defined in Equation 8, while *ê*_*ij*_ represents the importance score of the sampled edge, calculated using the node embeddings, as detailed in the original PGExplainer model.

We selected several correctly predicted DNA-binding sites from the protein sequences listed in Supplementary Table 1 to analyze the interpretability of both the sequence-based and structure-based models. The results are presented in Figure 5. The graph interpretability generation and visualization were implemented according to Lars et al. [41] using the PyTorch Geometric [42] package. As shown in Figure 5, the global attention mechanism allows the target site to capture long-range dependencies within the sequence. For example, residue 6 in protein 5D8C, chain B, shows high attention weights for residues around positions 20 and 40 (Figure 5A), while residue 51 in protein 6EN8, chain A, interacts strongly with residues around positions 10 and 30 (Figure 5B). Furthermore, attention weights are generally moderate when the residues are distant from the target site, which highlights the inherent advantage of attention-based models, such as Transformers, over recursive neural networks (RNNs). This phenomenon has been well-documented in related studies [43, 26].

**Figure 5:**
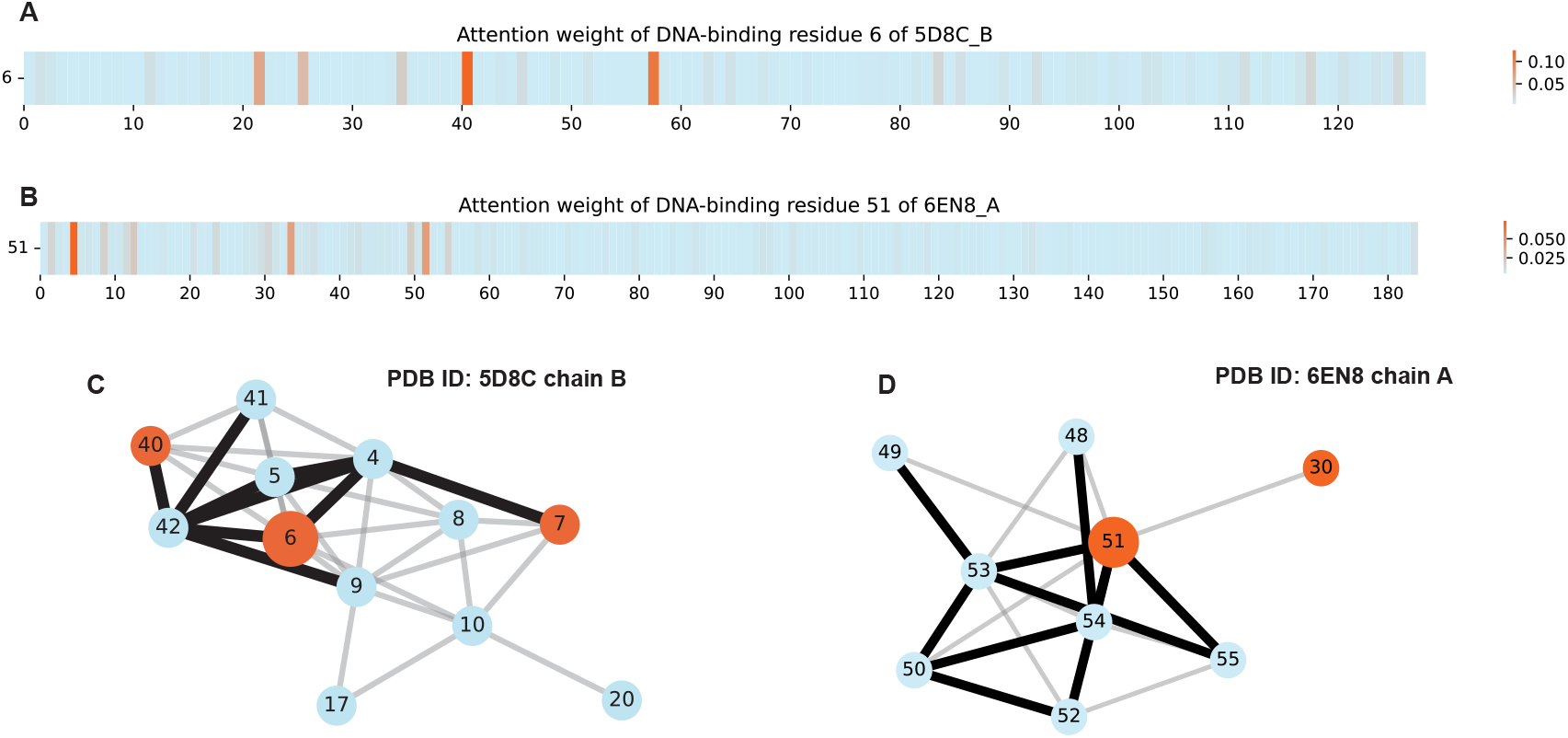
Interpretability analysis of the sequence-based and structure-based models. (A) Attention weight visualization for DNA-binding site 6 in protein 5D8C, chain B. (B) Attention weight visualization for DNA-binding site 51 in protein 6EN8, chain A. (C) Edge weight importance visualization for DNA-binding site 6 in protein 5D8C, chain B. (D) Edge weight importance visualization for DNA-binding site 51 in protein 6EN8, chain A. The edge weights are derived from the PGExplainer model. Red nodes represent DNA-binding residues, while blue nodes indicate non-binding sites. Bold edges highlight more important edge weights compared to the gray edges. The target residue is depicted with a larger node size.

It is widely recognized that large-scale attention-based pre-trained models can capture long-range interactions that reflect structural information within proteins. This conclusion has been drawn from downstream tasks such as classification using encoder blocks and generative predictions using decoder blocks [44, 45]. Our study reached similar conclusions through the interpretability analysis of a parallel structure-based model. As shown in Figure 5, the bold edges represent the top 10 edge weights in the 1-hop subgraph of the target residue. For instance, residue 6 in protein 5D8C, chain A, exhibits stronger attention to residues around positions 20 and 40. Residues 20, 40, 41, and 42 are the 1-hop neighbors of residue 6 (Figure 5C) and have high edge weights, indicating that this subgraph plays a crucial role in predicting residue 6 as a DNA-binding site. Interestingly, the attention weights between residue 6 and the residues around position 20 are smaller than those with residues around position 40, which is consistent with the edge weights in the subgraph. Additionally, the residues near position 20 form a cluster that contributes to graph prediction as 2-hop neighbors (Supplementary Figure 4A).

Similarly, in protein 6EN8, chain A, residue 51 shows larger edge weights with spatially proximate residues (Figure 5D). Notably, DNA-binding residue 30 is a 1-hop neighbor of residue 51, as reflected in both the attention weight visualization and the subgraph. The residues surrounding position 30 also form a 2-hop subgraph cluster (Supplementary Figure 4B).

In conclusion, our analysis of both the sequence-based and structure-based models using attention scores and edge weights revealed that the attention-based model successfully learned structural information from a novel interpretability perspective. These results suggest that pre-trained language models capture sufficient information to make accurate predictions, particularly in cases where protein structure is unavailable. This interpretability analysis also helps to explain the comparable performance of sequence-based and structure-based models, as demonstrated in Section 3.1.

## 4 Discussion

Protein-DNA binding plays a crucial role in numerous biological processes, and accurately identifying the DNA-binding residues of a given protein is essential as the first step in modeling protein-DNA interactions. This is particularly important for downstream tasks such as motif analysis [46] and drug design [47]. Although deep learning techniques have led to the development of several models that rely on either protein sequence or structural information, obtaining high-resolution protein structures remains both time-consuming and expensive. As a result, the number of solved DNA-binding protein structures represents only a small proportion of those without available structures. Despite the substantial advances made by AlphaFold2 [34], the performance of DNA-binding prediction models is still largely influenced by the accuracy of predicted structures [12]. Thus, there remains a pressing need to develop accurate sequence-based models for DNA-binding residue prediction using only sequence data.

In our study, we capitalized on the strengths of large-scale pre-trained protein language models and attention mechanisms to develop a sequence-based model. Additionally, we incorporated an effective contrastive learning module to enhance the model’s ability to learn discriminative features in the embedding space. We evaluated our model on two widely used benchmark datasets, and as shown in the results, our model outperformed all existing sequence-based models (Figure 2, Table 2) and demonstrated strong generalization capabilities on unseen datasets. Our analysis suggests two primary reasons for this superior performance: (1) the pre-trained model effectively extracts sequence information from a large-scale dataset of protein sequences, which is sufficient for subsequent tasks [48, 49]; and (2) the attention-based model captures long-range dependencies within the protein sequence more efficiently than the convolutional models employed by the second-best model, DBPred [8].

We also applied a contrastive learning module, inspired by Wang et al. [33], and designed a structure-based model using a graph convolutional network (GCN) to validate that the contrastive learning module can serve as a general component in the DNA-binding prediction task. While the contrastive learning module made only limited improvements to the sequence-based model, the t-SNE dimension reduction visualizations for both sequence-based and structure-based models (Figure 3) revealed that residues in the structure-based model are more distinguishable than those in the sequence-based model. This indicates that the attention-based model already captures sufficient information, limiting the performance gains from contrastive learning. However, when we replaced the attention-based module with a multi-layer perceptron (MLP), the contrastive learning module led to a significant improvement in performance, suggesting that it is particularly effective when applied to simpler architectures. As our experiments further showed (Figure 4), model performance can increase when the contrastive learning module is applied to larger embedding dimensions, although the appropriate dimension size must be determined experimentally. In summary, contrastive learning can be viewed as a flexible component, particularly useful for handling imbalanced datasets, with its location and embedding size needing to be adjusted based on model complexity.

In Section 3.3, we performed an interpretability analysis and demonstrated that the sequence-based model learned structural information from a novel perspective. By analyzing the attention scores of the sequence-based model and the edge weights from the structure-based model, we observed overlaps between regions with high attention scores and regions that are spatially proximate in protein structures (Figure 5). Our interpretability analysis suggests that the attention mechanism tends to focus on spatially adjacent residues, even in the absence of structural information. Additionally, the edge weights of the structure-based model were found to be higher for directly connected neighboring nodes, which could be a result of the cutoff value *c* in Equation 7. This cutoff limits connections to nodes within a certain distance, potentially leading to over-smoothing issues [50] when information is passed from distant nodes in deep GCN models. Recent studies have explored using Transformer models to process graph-like structures with fully connected graphs [51, 52], and our findings suggest that the global attention mechanism could be directly interpreted as a fully connected graph.

In conclusion, we developed an attention-based deep learning model for predicting DNA-binding residues using only protein sequence information. By utilizing a large-scale pre-trained model as a feature extractor without fine-tuning and incorporating a contrastive learning module for learning discriminative embeddings, our model achieved state-of-the-art performance among existing sequence-based models. Furthermore, our interpretability analysis revealed that the sequence-based model successfully captured structural information, offering a novel perspective on model interpretability in the field.

## TABLES

**Supplementary Table 1:** Model performance of AUC per sequence.

| PDB ID | Chain | Sequence-based | Structure-based | Difference |
| --- | --- | --- | --- | --- |
| 6EN8 | A | 0.993 | 0.983 | 0.010 |
| 5VXN | C | 0.985 | 0.975 | 0.010 |
| 5JUB | A | 0.972 | 0.982 | 0.010 |
| 6EE8 | M | 0.963 | 0.971 | 0.008 |
| 5VL9 | A | 0.961 | 0.968 | 0.007 |
| 5F7Q | E | 0.960 | 0.961 | 0.001 |
| 5D8C | B | 0.955 | 0.945 | 0.010 |

---

**Supplementary Figure 1:**
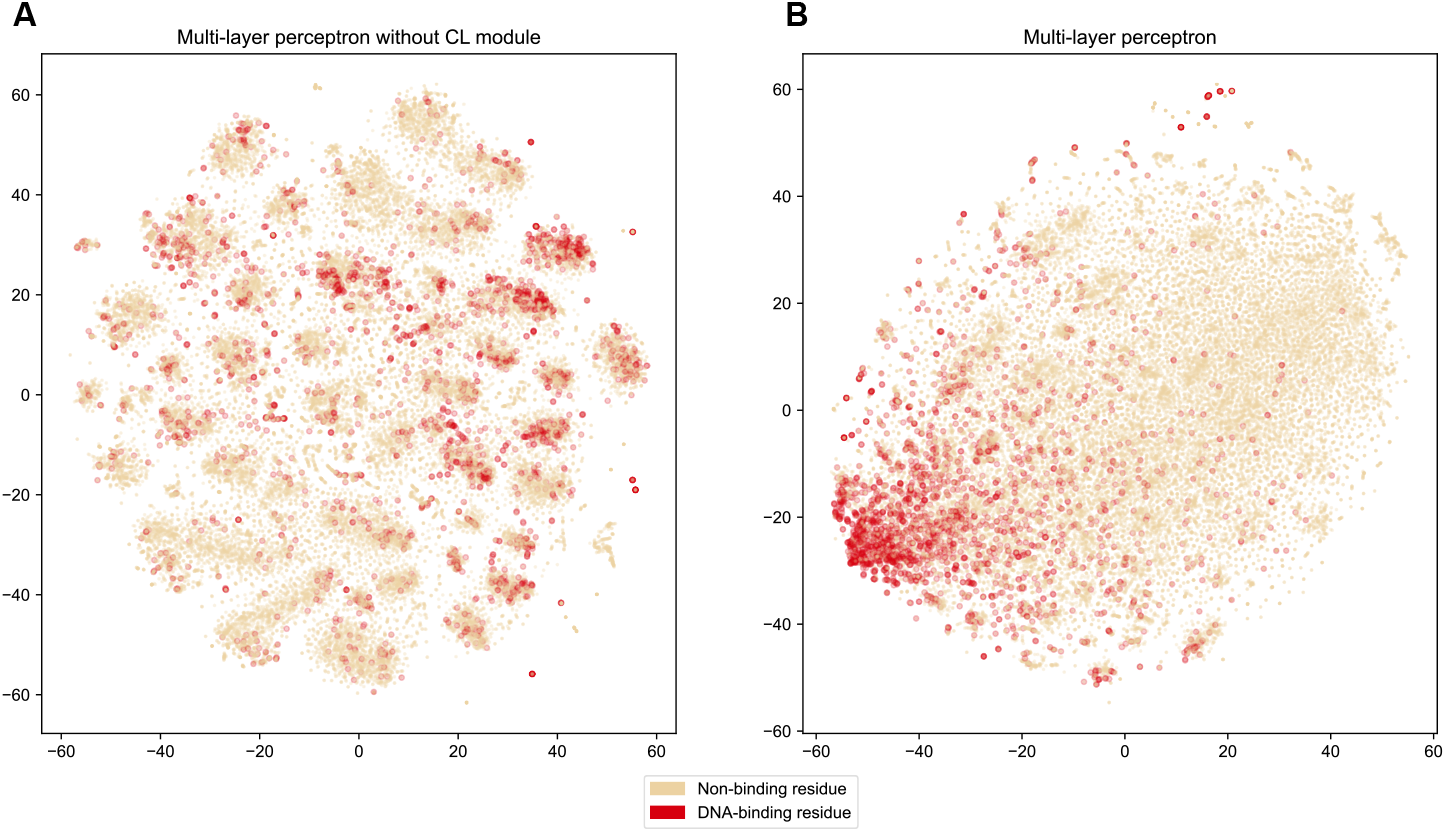
The t-SNE reduction visualization of the embedding space for the multi-layer perceptron (MLP) model tested on TE129. (A) Embedding space of the MLP model without the contrastive learning module. (B) Embedding space of the MLP model with the contrastive learning module.

**Supplementary Figure 2:**
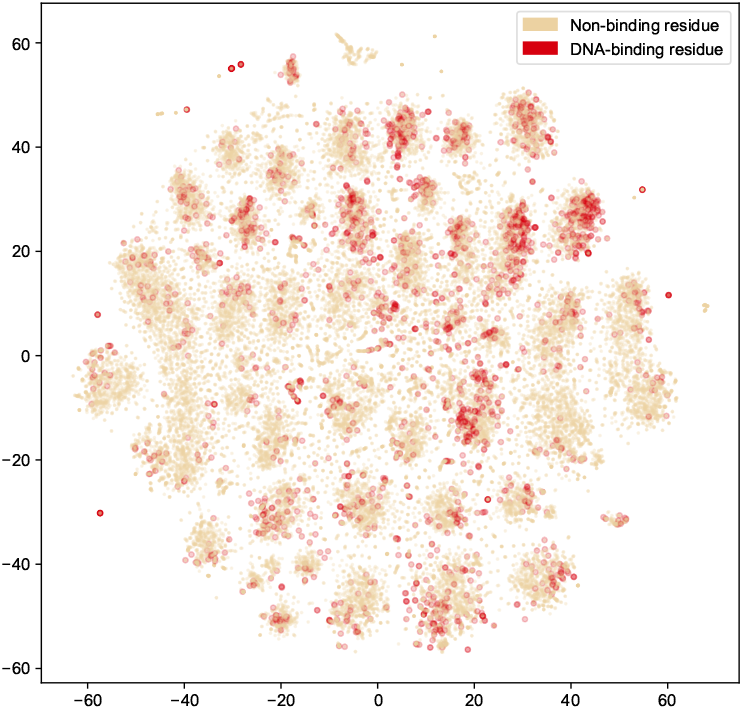
The t-SNE reduction visualization of the initial embedding space of TE129, generated by ProtBert.

**Supplementary Figure 3:**
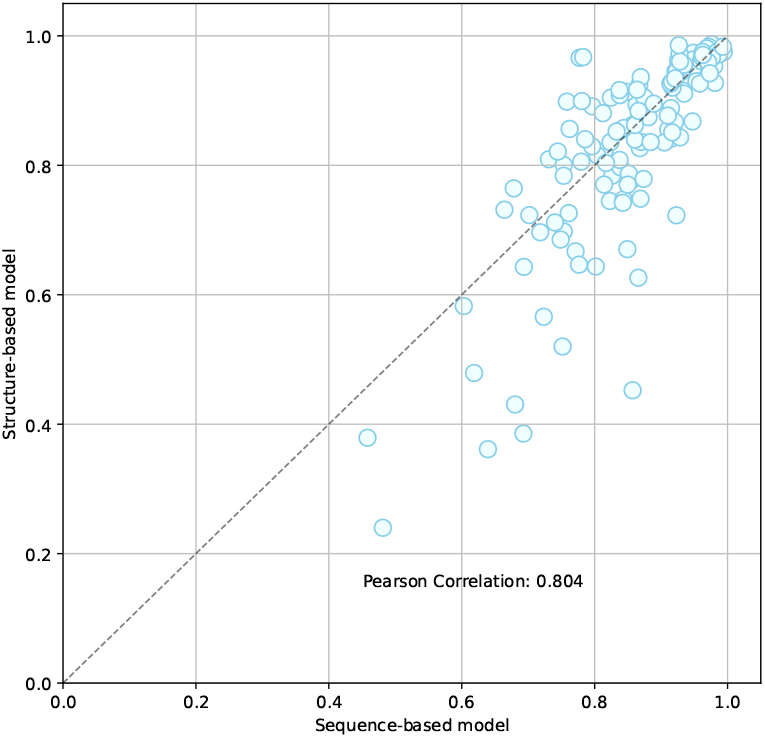
The correlation of AUC per sequence between the sequence-based and structure-based models on TE129.

**Supplementary Figure 4:**
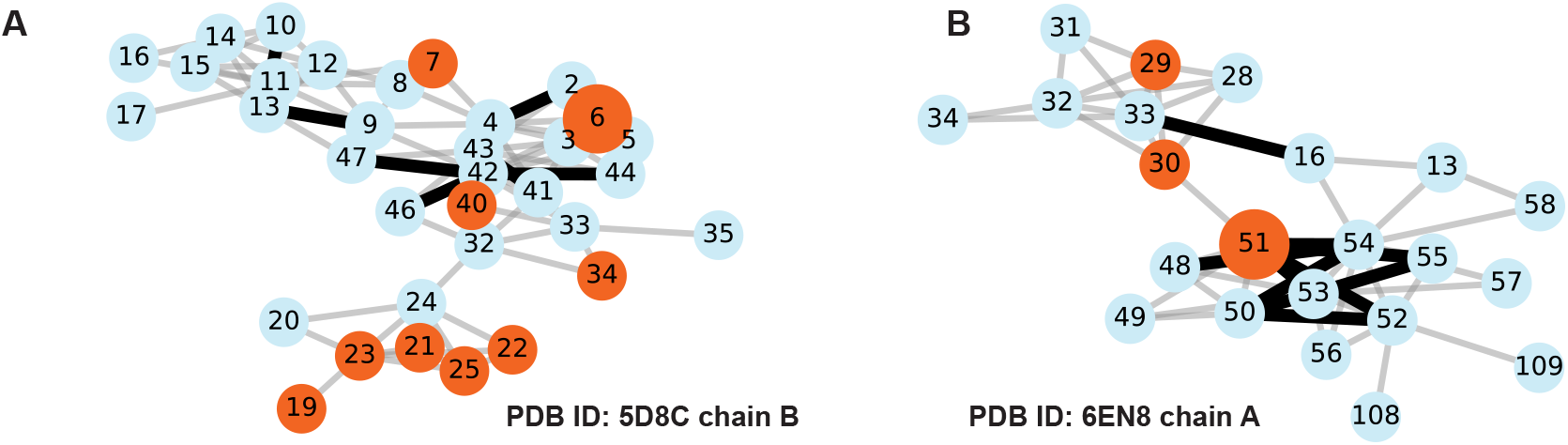
The visualization plot of the 2-hop subgraph for the target residue in (A) protein 5D8C, chain B, and (B) protein 6EN8, chain A.

